# Women’s health outreach using a rapid and fieldable Nanopore-based microbiome profiling workflow

**DOI:** 10.64898/2026.11.09.765251

**Authors:** Sander Boden, Chenay Waldekker, Margaretha E.H. Kaijen-Lambers, Nicole P.W.C.J. van den Braak, Daan C.J. Brackel, Linda Wijnstroom-de Quartel, Luena Pinheiro Keulers, Eline S. Klaassens, Cindy M.A. de Bot, Margriet A. van Gestel, Eefje J.A. Schrauwen

## Abstract

**Background:** In recent years, scientific interest in the vaginal microbiome has increased due to its impact on women’s health. Colonization of the *Lactobacillus* species in the vagina may contribute to protection against vaginal infectious diseases. In order to create public awareness regarding vaginal health and to provide insights into the vaginal microbiome, a rapid field microbiome analysis using Nanopore sequencing was evaluated at Lowlands, a three-day music festival in the Netherlands. This study aimed not only to evaluate a rapid sequencing workflow, but also to explore citizen science as a tool for health promotion and public engagement in women’s health.

**Methods:** The workflow at Lowlands contained the following steps; (i) Dialogue concerning vaginal health, with invitation to participate. (ii) Informed consent and questionnaire to investigate knowledge on vaginal health, life-style and contraceptive usage; (iii) self-collection of a vaginal swab; (iv) DNA isolation using QuickExtract DNA extraction solution; (v) Ultrafast amplification of the 16S rRNA gene using NextGenPCR™ thermal cycling using 16S Barcoding kit 24 V14; (vi) Library preparation and sequencing was performed according to the manufacturers protocol using R10 chemistry; (vii) For data-analysis, MABA16S was used, with slight modifications. This workflow was established within 4 hours and results were communicated with interested participants.

**Results:** A total of n=303 people participated in this study within a timeframe of three days, with n=265 samples being analyzed. 35.8% of participants were dominated by *L. crispatus*, followed by *L. iners* (26.4%) and *L. jensenii* (6,4%). Additionally, lifestyle characteristics for *Lactobacillus* species carriership were investigated, with a statistically significant association found between dominance of *Lactobacillus* species and the contraceptive used. A knowledge gap was found regarding the awareness of *Lactobacillus* species in the vagina. Also, the project stimulated dialogue about vaginal health, with participants reporting that the study increased their awareness and ability to talk openly about the topic.

**Conclusions:** This project successfully developed a rapid microbiome analysis capable of mapping the vaginal microbiome within four hours. Additionally, the findings highlight that citizen science-based research can effectively spark important conversations and may help break the taboo surrounding vaginal health.

**Clinical trial number:** not applicable.

## Background

In recent years, there has been a growing scientific interest in the vaginal microbiome due to its impact on female health. *Lactobacillus* species typically dominate 70% of the human vaginal microbiome of women of reproductive age (1). These *Lactobacillus* species, specifically *L. crispatus*, are crucial in their interaction with the host’s innate immune system, and research shows that a low abundance of *Lactobacillus* increases the risk of various vaginal infections (2). Hormonal contraception and hygiene can support the balance of the microbiome, while other factors can disrupt it, increasing risks of bacterial vaginosis and sexually transmitted infections (STIs) (3). Since daily habits and lifestyle can influence the vaginal ecosystem, promoting a healthy vagina is crucial.

The general public lacks knowledge about vaginal health and the microbiome associated with it (4). Normalizing the conversation about vaginal health is crucial for wellness and prevention, striving for a healthier future for all women. Collaborating with citizen scientists is a powerful way to achieve this goal (5–7). Citizen science enables researchers to conduct large-scale data collection while fostering knowledge transfer to the public, breaking down taboos, and involving participants in the analysis of results. A practical course provides an accessible way for individuals to gain a better understanding of their own bodies by studying their own samples. In such a setting, knowledge is directly transferred, taboos can be more easily addressed, questions can be raised, and science is brought closer to the individual, which has already proven effective on a smaller scale in conveying knowledge about vaginal health (5).

Citizen science has increasingly been recognized as a valuable approach within public health research, as it allows citizens to actively participate in data collection, interpretation, and knowledge dissemination. This participatory model not only broadens the scope and diversity of scientific datasets but also strengthens health literacy and empowers individuals to engage with their own health (5,8). In the field of women’s health, citizen science has proven particularly powerful for addressing sensitive topics such as the vaginal microbiome, where stigma and lack of awareness often hinder open dialogue (3,4). By involving participants in both the sampling process and the discussion of results, research initiatives can contribute simultaneously to scientific knowledge and to the normalization of conversations around reproductive health. This dual function positions citizen science as not merely a methodological tool, but also as a public health intervention with the potential to improve awareness, reduce taboos, and foster preventive health behavior (3,8).

In order to actively involve the public in the analysis of results, an ultra-fast diagnostic method was developed to screen vaginal swabs for the presence of *Lactobacillus* species; this had never before been achieved within a time frame of four hours and in a field-laboratory setting. The development of such rapid and field-deployable assays is of great importance for the future, as they enable microbiome studies to be performed outside conventional laboratory environments. A method for fast, fieldable microbiome assessment can be achieved through a combination of rapid nucleic acid extraction, ultrafast PCR, and portable sequencing technologies, opening new possibilities for point-of-care diagnostics and large-scale citizen science applications. This concept inspired the test-case conducted at the Lowlands science site. Within that test case, we investigated which factors are significant predictors of *Lactobacillus* species carriage. Through this research, we also aim to break the taboo surrounding vaginal health by encouraging open dialogue.

## Methods

### Study cohort and data collection

A field lab study was conducted from August 16, 2024 to August 18, 2024 at Lowlands science site, Biddinghuizen, The Netherlands. Inclusion criteria were an age of 18-45 years, with no active pregnancy or menstruation at the time of sampling. A total of 303 vaginal samples were collected using a sterile FLOQS swab (Copan) and Liquid Amies Transport Medium (52000, Molecular Biology Systems, The Netherlands). After obtaining an informed consent, the participants were invited to complete a short survey using the platform Qualtrics [see Additional file 1]. No personally identifiable information was gathered. One vaginal swab was self-collected in a standardized way by the participants, with sampling rooms containing instructions in layman terms. Immediately after sampling, swabs were transferred to a vial and stored at 4°C until further processing. The subset of samples that could not be processed on-site were kept at 4°C until further processing.

### DNA isolation

DNA was isolated using the QuickExtract DNA extraction solution (QE09050, ImmunoSource, Belgium) following the manufacturer’s protocol with slight modifications. Specifically, 400 µl of sample was used and 6 µl of 50 mg/mL lysozyme (L7651, Sigma-Aldrich, USA) was added, followed by incubation at 37°C and 400 rpm for 2.5 minutes. Afterwards, 500 µl of QuickExtract solution was added, followed by 15 seconds of vortexing, 6 minute incubation at 65°C, 15 seconds of vortexing and a final 2 minute incubation at 98°C. The resulting solution was directly used as template for the PCR reaction. As a control the zymoBIOMICS microbial Community Standard (D6300, Zymo research, CA) was used.

### NextGenPCR™ and 16S rRNA gene sequencing

For amplification and sequencing of the full-length 16S rRNA gene (∼1500bp), the 16S barcoding kit 24 V14 (SQK-16S114-24, Oxford Nanopore Technologies, UK) was adapted for use with the NextGenPCR™ Thermal Cycler (10001; Molecular Biology Systems, the Netherlands). For the PCR master mix, 2x Arctic Fox HF Chemistry-2x (50050-1, Molecular Biology Systems, the Netherlands), 1% DMSO, 2 µl template DNA and 6.5 µl barcoded primer (SQK-16S114-24, Oxford Nanopore Technologies, UK) were used with a final volume of 20 µl. The following PCR conditions were used: initial denaturation at 98 °C for 60 s, 40 cycles of 98 °C for 10 s, 57 °C for 30 s, 68 °C for 10 s, and a final extension step of 68 °C for 60 s. The PCR product concentrations were measured using the Qubit 3.0 fluorometer (Q33216, Thermo Fisher Scientific, US) with the dsDNA HS Assay Kit (Q32851, Thermo Fisher Scientific, US). All further processes followed the manufacturer’s instructions. In order to limit turnaround time, sequencing was performed for 30-40 minutes, aiming for >1k reads per sample.

### Data analysis

The quality of the sequencing data was assessed with MinKNOW V24.02.16 (default settings). Samples with <500 reads or (partially) unanswered surveys were excluded from further analysis. The 16S rRNA gene sequencing data was analyzed using MABA16S v0.1.0 using default settings (9), and the results were subjected to an abundance threshold of >1%. Classified taxa with abundances <1% were grouped and labeled as ‘other’. Additional post-processing was performed by excluding samples that contained <200 classified reads. Application of this threshold ensures the detection of a dominant species (>51%) is supported by at least 100 reads. Before statistical analysis, the abundances were converted to a binary table describing the dominant species (>50% abundance), if any were present.

### Statistical analysis of the association between lactobacillus dominance and characteristics

To assess associations between various categorical variables (age group, menstrual products, hygiene products, contraceptive methods, and sexual behavior) and the presence of dominant species of *Lactobacillus*, chi-square tests of independence were performed.

Prior to analysis, the data were cleaned and categorical variables were one-hot encoded to ensure proper handling. For menstrual and hygiene product variables, where respondents could select multiple options, separate binary variables were created for each product, indicating whether a participant used that product (1) or not (0).

For each analysis, contingency tables were created between the categorical variable of interest and the *Lactobacillus* species: dominant *Lactobacillus* species, dominant non-*Lactobacillus* species or presence of *Lactobacilllus crispatus*. Categories with very low counts or unclear labels (e.g. “*L. gallinarum*”, “Other, namely”, and “I am sterilized”) were excluded from the analyses, as they did not provide sufficient data for meaningful comparison. These subgroups are, however, displayed in descriptive figures for completeness.

Chi-square tests were conducted using Monte Carlo simulation to estimate p-values with 10,000 replicates (simulate.p.value = TRUE, B = 10000), improving the accuracy of the p-value estimation in cases of small or unevenly distributed samples, as recommended by Bradley & Cutcomb (10). This approach was implemented using the chisq.test function in R.

## Results

### Scientific outreach and knowledge transfer during Lowlands

During Lowlands, numerous conversations were engaged with a diverse group of individuals. Many women expressed their appreciation for the research, emphasizing the need for an open dialogue about the vagina. They reported feeling empowered to share their stories and ask questions about discomforts and concerns. We observed that conducting research in collaboration with citizens effectively initiates important conversations. Not only did women express their interests, but also men showed interest in learning more about the vaginal microbiome. They noted that gaining knowledge on this topic would enable them to discuss it more comfortably with their partners. Interestingly, only 17% of the participants in this study were familiar with the *Lactobacillus* genus.

### Lowlands fieldlab inclusion & population descriptors

A total of 303 participants aged <21-45 years signed up to participate in the research. In total, 265 samples met the inclusion criteria and were used for further analysis (Figure 1). Among the included participants, the majority were aged between 21 and 40 years old (n=256) [see Additional file 2]. The smallest sample sizes were found in the group younger than 21 (n=3) and aged 41-45 years (n=6). Among hygiene products, the least used was the vaginal douche (n=1) and water was the most commonly used (n=240). During the field study, 48 samples were sequenced per day, with these participants receiving their results (presence/absence of *Lactobacillus*) at the end of each day. The remaining samples that met the inclusion criteria were sequenced at Avans University of Applied Science, Breda.

**Figure 1:**
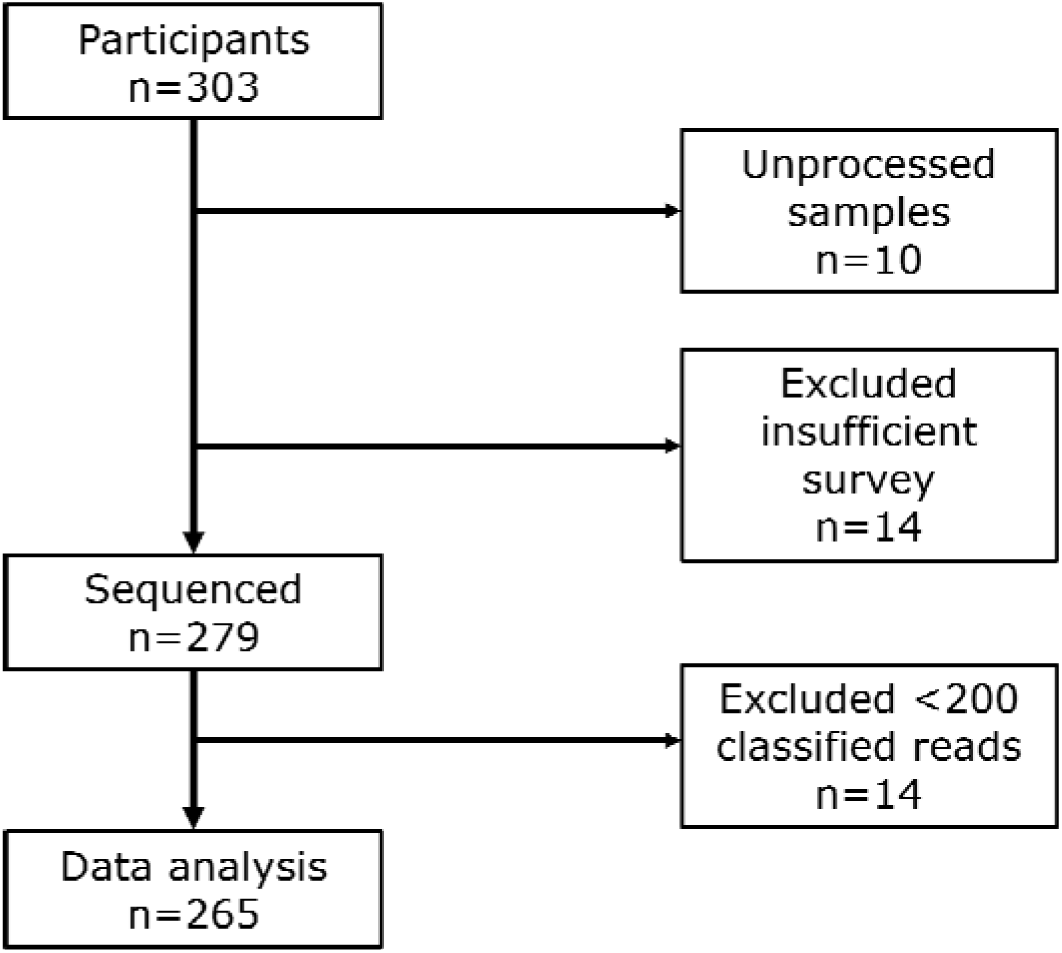
Vaginal swab samples excluded from analysis, split by reason.

### Lowlands vaginome composition

After taxonomic classification of the sequencing data, dominant taxa were determined for each sample. The majority (71.3%, n=189) of participants’ microbiomes were dominated by one of *Lactobacillus crispatus* (35.8%, n=95), *Lactobacillus iners* (26.4%, n=70), *Lactobacillus jensenii* (6.4%, n=17), *Lactobacillus gasseri* (2.3%, n=6), or *Lactobacillus gallinarum* (0.4%, n=1). Among the remaining 28.7% (n=76) of participants, 63.2% (n=48) had a non-*Lactobacillus* dominant species in their vaginal microbiome, and the remaining 36.8% (n=28) had no dominant taxon present, which were classified as variable microbiomes (Figure 2). No co-dominance of any two taxa was observed among the samples without a dominant taxon.

**Figure 2:**
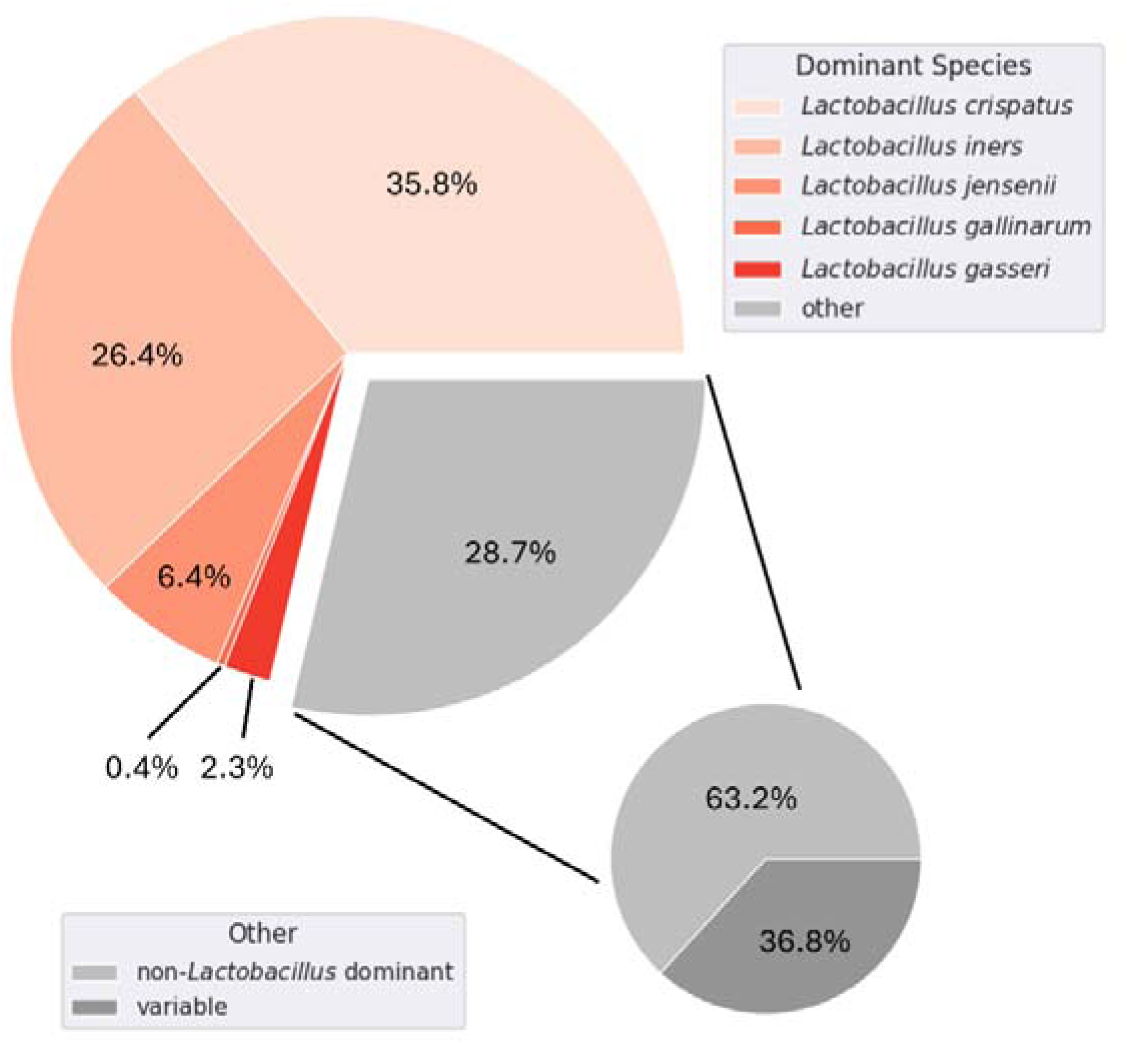
Dominant species found in the vaginal microbiomes of Lowlands participants.

### Determinants of *Lactobacillus* carriage

In order to evaluate the effect of lifestyle and intimate hygiene on dominance of *L. crispatus* or any *Lactobacillus* species, participants answered a short questionnaire about their age, use of menstruation products and intimate hygiene products, recent drug use, unprotected sexual activity and contraceptive use. Chi-square tests for dominance of *Lactobacillus* species or *Lactobacillus crispatus* were performed for each question [see Additional file 2]. Of all the determinants included in the questionnaire, only the type of contraception used by participants resulted in a significant association with the dominance of a *Lactobacillus* species (36.6; p < 0.01), and more specifically, *L. crispatus* (12.3; p < 0.01). Post-hoc pairwise comparisons between specific contraceptive methods (oral birth control, copper IUD, hormonal IUD, sterilization, other, or no contraception) using Bonferroni-Holm correction showed a significant difference between the usage of hormonal IUD or birth control pills and no contraceptive use (Figure 3). Notably, the percentage of *L. crispatus* dominant samples was higher among participants who used either birth control pills (n=44; 48%) or hormonal IUD’s (n=101; 41%), when compared to participants who did not use contraception (n=84; 25%).

**Figure 3:**
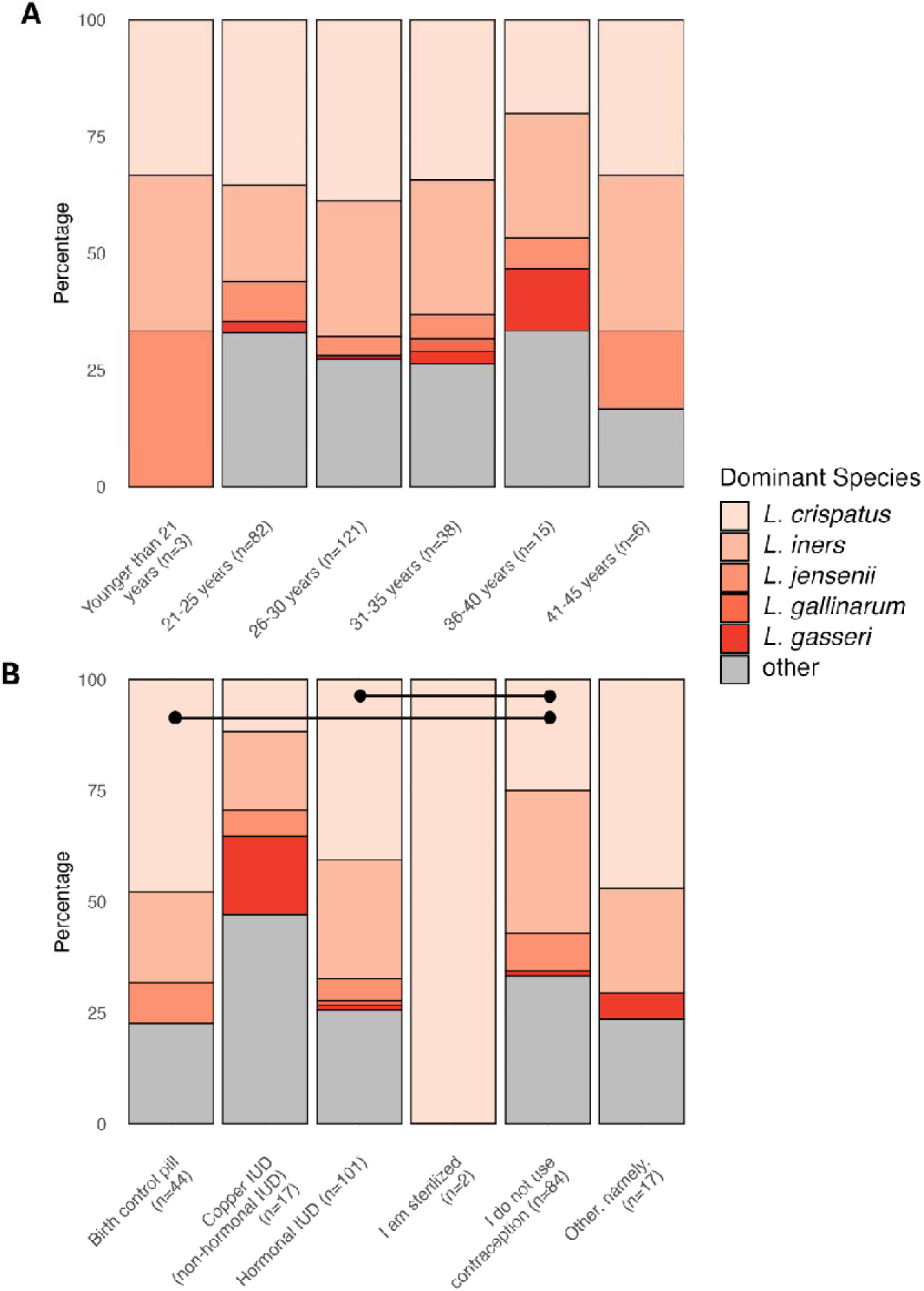
*Lactobacillus* species dominance within groups of age (A) and users of various contraceptive methods (B). Horizontal lines indicate significant differences between groups after Bonferroni correction (p < 0.05).

## Discussion

In this study, we present a rapid microbiome analysis which can be applied in a field lab setting with the aim of breaking the taboo around vaginal health by promoting open dialogue while collecting valuable data. In addition, we examined features such as age, menstrual products, and hygiene product used to determine if they significantly predicted *L. crispatus* carriage. Through optimization of existing 16S rRNA gene sequencing protocols, and adaptation to the faster NextGenPCR™ Thermal Cycler, we developed a rapid, light-weight workflow for investigating the vaginal microbiome in a field-setting. This workflow enabled the determination of dominant bacteria in the vaginal microbiome within four hours.

Using this workflow, the “Lowlands microbiomes” were found to be mostly dominated by *L. crispatus* (35.8%), followed by *L. iners* (26.4%). Notably, among 28.7% of participating individuals, the microbiome was either variable (36.8%) or dominated by a non-Lactobacillus species (63.2%). These findings are comparable to a Belgian study from 2023 where, within a cohort of 3000+ individuals, 43% of microbiomes were found to be dominated by *L. crispatus*, followed by 28% dominated by *L. iners* (3). Interestingly, the authors also reported 10% of microbiomes to be dominated by species of the *Gardnerella* genus, which was not found among the cohort presented in this study. One sample classified as a variable microbiome was found to contain a small percentage (2%; 53 reads) of *Gardnerella vaginalis*, indicating the ability of our data-analysis workflow to classify these species. Previous research has shown primer-specific biases in 16S sequencing results, which likely explains the discrepancy in *Gardnerella* abundances (11,12).

While our workflow allowed rapid and on-site 16S rRNA gene sequencing and subsequent data analysis, limitations are present in the due to time-constraints and processing capacity. In order to report results to the participants on the same day, concessions were made with regards to sequencing depth. With this limited sequencing depth (>500 reads, >200 classified reads), the choice was made to only analyze and report the dominant taxa within the microbiome to the participants. Accurate determination of microbial composition requires a higher sequencing depth that could not be achieved while balancing the other goals of our presented study (13). More specifically, the vaginal microbiome is complex, where co-dominance of certain species is often seen, along with more variable and highly diverse microbiomes (3). In addition, sampling was performed at Lowlands, a Dutch music festival, which may be prone to sampling bias.

Statistical analysis into the predictive power of lifestyle choices for the dominance of *Lactobacillus* species and *L. crispatus* showed a significant association for contraceptive use. Post-hoc analysis indicates that the use of birth control pills and hormonal IUDs significantly increases the chance of a microbiome dominated by *L. crispatus* when compared to no contraceptive use [see Additional file 2]. Current research reports higher prevalence of bacterial vaginosis and dysbiosis in copper IUD users whereas hormonal contraceptives are considered to provide a stable environment for healthy vaginal microbiomes (14–16). While these findings are still conflicting and contradictory, our results support the effects of hormonal contraceptives on the vaginal microbiome.

Beyond the methodological innovation of rapid sequencing in a field-laboratory setting, this study illustrates how citizen science can function as a public health strategy. By directly involving participants in the collection and interpretation of their own vaginal microbiome data, we created an accessible setting where health information was personalized, understandable, and immediately relevant. Such approaches can support the public health goals of prevention and early awareness, by improving health literacy and enabling individuals to recognize the importance of maintaining a healthy vaginal microbiome (7,8).

Our observations at Lowlands show that citizen science can help to overcome stigma, generate open dialogue, and reach groups who may not normally be exposed to this type of health education. These outcomes align with public health objectives that emphasize empowerment, equity, and community-based approaches to health promotion (5). Moreover, by embedding science in a non-clinical and informal environment, barriers to participation were lowered and the topic of vaginal health became approachable, which is crucial for breaking taboos and stimulating preventive behavior (4).

Citizen science projects such as this not only enrich scientific understanding of the vaginal microbiome but also serve as innovative public health interventions. Scaling up such initiatives to schools, community centers, or primary care could provide new opportunities to integrate microbiome knowledge into everyday health promotion. In this way, citizen science can bridge the gap between research and practice, contributing to collective responsibility for women’s health and advancing broader public health agendas.

Vaginal health remains a critical topic warranting further research. Inclusion of citizen scientists in this research is a way to open the conversation and break taboos surrounding the subject of vaginal/women’s health. In future efforts, small-scale citizen science projects could educate people on this matter.

In conclusion, the vaginal microbiome is a complex and dynamic system influenced by various factors such as menstrual products, contraceptive methods, and the natural hormonal fluctuations associated with age and the menstrual cycle. While research shows that products like menstrual cups and hormonal contraceptives may support a healthier microbiome by promoting lactobacillus dominance, conflicting evidence regarding non-hormonal contraceptive methods such as copper IUDs underscores the need for more comprehensive studies. Understanding how these elements interact with the vaginal microbiome is critical for improving women’s reproductive health. Future research, including longitudinal studies and citizen science initiatives, can help clarify these relationships and empower women to make informed choices about their reproductive health.

## Additional files

**Filename:** Additional file 1.pdf

**Title:** Survey for collecting lifestyle characteristics.

**Description:** The survey used during sample collection to collect participants’ lifestyle characteristics.

**Filename:** Additional file 2.pdf

**Title:** Statistical analysis of determinants.

**Description:** A table containing dominance percentages and the statistical associations of *Lactobacillus* species and *L. crispatus*, grouped by determinants.

## List of abbreviations

IUD: Intrauterine device
PCR: Polymerase Chain Reaction
STI: Sexually transmitted infection

## Declarations

### Ethics approval and consent to participate

This study was approved by the Ethical Committee of Avans University of Applied Science (24.008), in accordance with the Dutch code for scientific integrity (NGWI). Informed consent to participate was obtained from all participants.

### Consent for publication

Not applicable.

### Availability of data and materials

The accession referenced in the manuscript (PRJEB106352) is available through the European Nucleotide Archive (ENA): https://www.ebi.ac.uk/ena/browser/view/PRJEB106352

### Authors’ contributions

Conceptualization, E.J.A.S.; Formal analysis, S.B., C.W., M.A.G., E.J.A.S.; Investigation, S.B., C.W., M.E.H.K.L., C.M.A.B., N.P.W.C.J.B., D.C.J., L.W.Q., L.P.K., E.S.K., M.A.G., E.J.A.S.; Methodology, S.B., C.W., E.J.A.S.; Project administration, E.J.A.S.; Resources, L.W.Q., E.J.A.S.; Supervision, E.J.A.S.; Visualization, S.B., M.A.G.; Writing—original draft, S.B., M.A.G., E.J.A.S.; Writing—review and editing, S.B., C.W., M.A.G., E.J.A.S. All authors have read and agreed to the published version of the manuscript.

## Acknowledgements

The authors thank Lowlands Science - organized by the Dutch Research Agenda, BKB and New Scientist - who provided the opportunity for this field-lab, and the participants at Lowlands for their willingness and enthusiasm for research into women’s health and the vaginal microbiome. We are grateful towards Yoni and Molecular Biology Systems for their participation with personnel and materials. Finally, we would like to thank Remco Kort, Shardelice Illidge and Rosanne Hertzberger for their fruitful discussions while initiating and performing this research.

