## Additional File 2 for "Women’s health outreach using a rapid and fieldable Nanopore-based microbiome profiling workflow"

| Determinants | | Sample size (n=265) | *Lactobacillus sp.* dominant | *L. crispatus* dominant | *Lactobacillus sp.* Dominance | | *L. crispatus* Dominance | |
|---|---|---|---|---|---|---|---|---|
| | | | | | Chi$^2$ | p | Chi$^2$ | p |
| Age | <21 years | 3 | 100% | 33% | 20.23 | 0.4227 | 2.187 | 0.8457 |
| | 21-25 years | 82 | 67% | 35% | | | | |
| | 26-30 years | 121 | 73% | 39% | | | | |
| | 31-35 years | 38 | 74% | 34% | | | | |
| | 36-40 years | 15 | 67% | 20% | | | | |
| | 41-45 years | 6 | 83% | 33% | | | | |
| Menstrual product use (multiple answers) | Pads | 118 | 70% | 29% | 11.37 | 0.4905 | 6.026 | 0.1189 |
| | Tampons | 185 | 72% | 55% | | | | |
| | Menstrual cup | 57 | 68% | 47% | | | | |
| | Menstrual underwear | 32 | 63% | 31% | | | | |
| | Other | 31 | 87% | 52% | | | | |
| Hygiene product use (multiple answers) | Humid tissues | 19 | 74% | 32% | 11.37 | 0.4925 | 2.960 | 0.4296 |
| | Soap/ showergel and water | 31 | 65% | 23% | | | | |
| | Lactacyd (washgel) | 6 | 83% | 50% | | | | |
| | Vaginal douche | 1 | 100% | 0% | | | | |
| | Water | 240 | 72% | 36% | | | | |
| Contraceptive use | No contraception | 84 | 67% | 25% | **36.65** | **0.0004** | **12.30** | **0.0057** |
| | Sterilization | 2 | 100% | 100% | | | | |
| | Hormonal intrauterine device (IUD) | 101 | 74% | 41% | | | | |
| | Copper intrauterine device (IUD) | 17 | 53% | 12% | | | | |
| | Contraceptive pill | 44 | 77% | 48% | | | | |
| | Other | 17 | 76% | 47% | | | | |
| Alcohol usage * | Yes | 249 | 72% | 37% | 3.263 | 0.4858 | 2.165 | 0.1851 |
| | No | 16 | 56% | 19% | | | | |
| Recreational drugs usage * | Yes | 103 | 73% | 38% | 2.909 | 0.5784 | 0.2975 | 0.6030 |
| | No | 162 | 70% | 35% | | | | |
| Unprotected sex * | Yes | 34 | 71% | 29% | 7.370 | 0.1150 | 0.7030 | 0.4475 |
| | No | 231 | 71% | 37% | | | | |

Note: rows marked with * refer only to the 24 hours preceding participation. Values in **bold** indicate significant ($p<0.05$) values.
