## Additional File 1 for "Women’s health outreach using a rapid and fieldable Nanopore-based microbiome profiling workflow"

**Informed consent for participation Viva La Vaginoom Lowlands 2024**

By participating in Viva La Vaginoom, you indicate that:

- You are at least 18 years old.
- You are not currently menstruating.
- You are not currently pregnant.
- You have not used excessive alcohol or drugs.

1. I have received sufficient information about this research project. The purpose of my participation as a participant in this study has been clearly explained to me, and I understand what participation entails.
2. My participation as a participant in this study is entirely voluntary. There is no explicit or implicit pressure for me to take part in this research.
3. It is clear to me that, if I object to one or more points mentioned above, I can stop my participation at any time, without giving a reason.
4. I give permission for the anonymous collection, storage, and use of my research results for answering the research question in this study.

I agree to participate in the Viva La Vaginoom study.
o Yes
o No

I give permission for reuse of my data after this study for currently unknown research within the field of (bio)medical education & research.
o Yes
o No

What is your assigned number? This is the number you received during registration.

What is your age group?
o Under 21
o 21–25
o 26–30
o 31–35
o 36–40
o 41–45

Are you familiar with the *Lactobacillus crispatus* bacterium?
o Yes
o No

Have you ever heard of community state types?
o Yes
o No

If you menstruate, which of the following products do you use during your menstruation? (Multiple answers possible)
o Sanitary pads
o Tampons
o Menstrual cup
o Menstrual underwear
o Other, namely

Do you use any of the following hygiene products?
o Lactacyd (wash gel)
o Wet wipes
o Vaginal douche
o Soap/shower gel and water
o Water only

How often do you use hygiene products? (per week)
• Frequency table from 0 to 7

Do you use any of the following contraceptive methods?
o Oral contraceptive pill
o Hormonal IUD (IUS)
o Copper IUD (non-hormonal IUS)
o I am sterilized
o I do not use contraception myself
o Other, namely

Have you had sex (without a condom) in the past 24 hours?
o Yes
o No

Have you consumed any of the following in the past 24 hours? (Multiple answers possible)
o Alcohol
o Drugs
o None of the above

Do you use an e-cigarette or vape?
o Yes, I vape
o Yes, I use an e-cigarette
o Yes, I use both
o No, I do not use an e-cigarette or vape

How often do you use an e-cigarette or vape? (per day)

- o Frequency table 0–30

Do you use another name for the word vagina?

Your responses have been recorded. On behalf of the entire Viva La Vaginoom team, we would like to thank you for your participation.
